# Stability and dynamics interrelations in a Lipase: Mutational and MD simulations based investigations

**DOI:** 10.1101/634253

**Authors:** Tushar Ranjan Moharana, Virendra Kumar, N. Madhusudhana Rao

## Abstract

Dynamics plays crucial role in the function and stability of proteins. Earlier studies have provided ambivalent nature of these interrelations. Epistatic effects of amino acid substitutions on dynamics are an interesting strategy to investigate such relations. In this study we investigated the interrelation between dynamics with that of stability and activity of *Bacillus subtilis* lipase (BSL) using experimental and molecular dynamics simulation (MDS) approaches. Earlier we have identified many stabilising mutations in BSL using directed evolution. In this study these stabilizing mutations were clustered based on their proximity in the sequence into four groups (CM1 to 4). Activity, thermal stability, protease stability and aggregations studies were performed on these four mutants, along with the wild type BSL, to conclude that the mutations in each region contributed additively to the overall stability of the enzyme without suppressing the activity. Root mean square fluctuation and amide bond squared order parameter analysis from MDS revealed that dynamics has increased for CM1, CM2 and CM3 compared to the wild type in the amino acid region 105 to 112 and for CM4 in the amino acid region 22 to 30. In all the mutants core regions dynamics remained unaltered, while the dynamics in the rigid outer region (RMSF <0.05 nm) has increased. Alteration in dynamics, took place both in the vicinity (CM2, 0.41 nm) as well as far away from the mutations (CM1, 2.6 nm; CM3 1.5 nm; CM4 1.7 nm). Our data suggests that enhanced dynamics in certain regions in a protein may actually improve stability.

**Statement of Significance:** How does a protein readjust its dynamics upon incorporation of an amino acid that improved its stability? Are the stabilizing effects of a substitution being local or non-local in nature? While there is an excellent documentation (from x-ray studies) of both local and non-local adjustments in interactions upon incorporation of a stabilizing mutations, the effect of these on the protein dynamics is less investigated. The stability and MD data presented here on four mutants, stabilized around four loop regions of a lipase, suggests that stabilizing effects of these mutations influence two specific regions leaving rest of the protein unperturbed. In addition, our data supports, observations by others, wherein enhancement in stability in a protein need not result in dampening of dynamics of a protein.

## Introduction

Proteins are flexible and rapidly fluctuating molecules with motions relevant to its function. Protein dynamics plays crucial role in their interaction with other macro- (protein, DNA, RNA etc.) and micro-molecules (ligand), enzymatic catalysis, protein transportation and protein degradation (1, 2). In particular, enzymatic catalysis, which requires stabilization of several intermediates and transition states, rely on protein dynamics to achieve the same (3). In fact, a direct correlation between enzyme dynamics and its activity has been established (4, 5). However, the relation between protein dynamics and its stability is more complicated. Both positive and negative correlations between stability and dynamics of enzymes have been reported (6–8).

Stable mutants often show compromised activity and vice versa (9, 10). The phenomenon is well documented and termed as activity-stability trade-off. It is speculated that such trade-off occurs via alteration of conformational dynamics (11, 12). Enzymes achieve enhanced stability by losing conformational dynamics, thereby leads to poor activity while hyperactive enzymes gain additional dynamics at the cost of stabilizing interactions. There is a common notion that enhanced conformational rigidity in the native state of the protein leads to increased thermostability. This hypothesis is supported by comparative analysis of dynamics of homologous proteins from psychro-, meso- and thermo-philic organisms (13) and laboratory evolved enzymes (14–17). However, deuterium exchange study of Rubredoxin from *Pyrococcus furiosus*, the most stable among the soluble protein known till date, at 28°C showed similar hydrogen-deuterium (H-D) exchange rate with that of other unstructured proteins (18). Berezovsky *et al.* have observed a statistical difference in arginine to lysine ratio in the proteomes of thermophiles compared to their mesophilic counterpart. By following all-atom Molecular dynamics simulation (MDS), they have shown that this observation is due to higher dynamics of arginine side chain than that of lysine in the native state of the proteins (13). Several other studies have been reported claiming a positive correlation between dynamics and stability of proteins (6, 19–22).

Different region in the protein vary in the magnitude of dynamics. An even more compelling question is how the dynamics of different region of an enzyme is altered by the mutations which increase enzyme stability without affecting its activity. To address this question we have combined mutations, having the above mentioned properties, in *Bacillus subtilis* lipase (BSL). Theses mutations were obtained earlier by our research group through methods of directed evolution. Stabilising mutations were combined based on their proximity in the BSL sequence to amplify the otherwise marginal effect of screened single mutants. Four combined mutants thus created, namely CM1, CM2, CM3 and CM4, were evaluated for their thermal stability, resistance to aggregation, resistance to protease hydrolysis and enzymatic activity along with BSL. Dynamics of BSL and mutants were studied by MDS. All the mutants showed improved thermal stability and resistance to aggregation and protease degradation over the wild type BSL, whereas two of them (CM2 and CM3) showed increased activity as well. Among the mutants, CM3 showed the maximum improvement both in terms of stability and activity. MDS has revealed that dynamics near two particular loops has increased in these mutants, irrespective of the position of mutations. In three of these four mutants, alteration of dynamics was observed to be taking place at a distance more than 1 nm from the site of mutations.

## Material

pET22b plasmid containing BSL gene was obtained from our previous work (23). Phusion master mix (2X) was used for PCR. All other enzyme and buffer used in molecular biology was purchased from NEB, India. *Para*-nitrophenyl butyrate (PNPB) and Triton X-100 was purchased from Sigma Aldrich, India. All other chemicals used are of at least analytical grade or better.

## Methods

### Creation of mutants

Stable yet active variants of BSL with multiple mutations were created by combining stabilizing single mutations obtained during our previous work (23). Mutations were clustered, based on their position in the protein sequence. Nearby mutations were combined by site-directed mutagenesis to generate stable mutants. In case of one position, wherein multiple stabilizing mutations were identified, the most stable variant was considered. Mutation I12A was ignored due to its poor activity. Four mutant lipases were purified and their secondary structure was recorded (Supplementary Fig.1). There are no discernible secondary structural differences between the mutants and the wild type lipase. Details of the mutations obtained and their properties are mentioned in the supplementary material. The above work resulted into four combined mutants (CM) containing 11 point mutations as described in table 1. The position of the mutations on the protein is shown in Fig. 1.

**Table 1:** Mutations in combined mutants. Mutations obtained from the screening were combined based on their sequence neighborhood.

| Mutant name | Mutations |
| --- | --- |
| CM1 | A15S, F17T |
| CM2 | T109V, G111S, L114P |
| CM3 | A132D, M134E, M137P |
| CM4 | G155S, I157M, Y161N |

**Fig.1:**
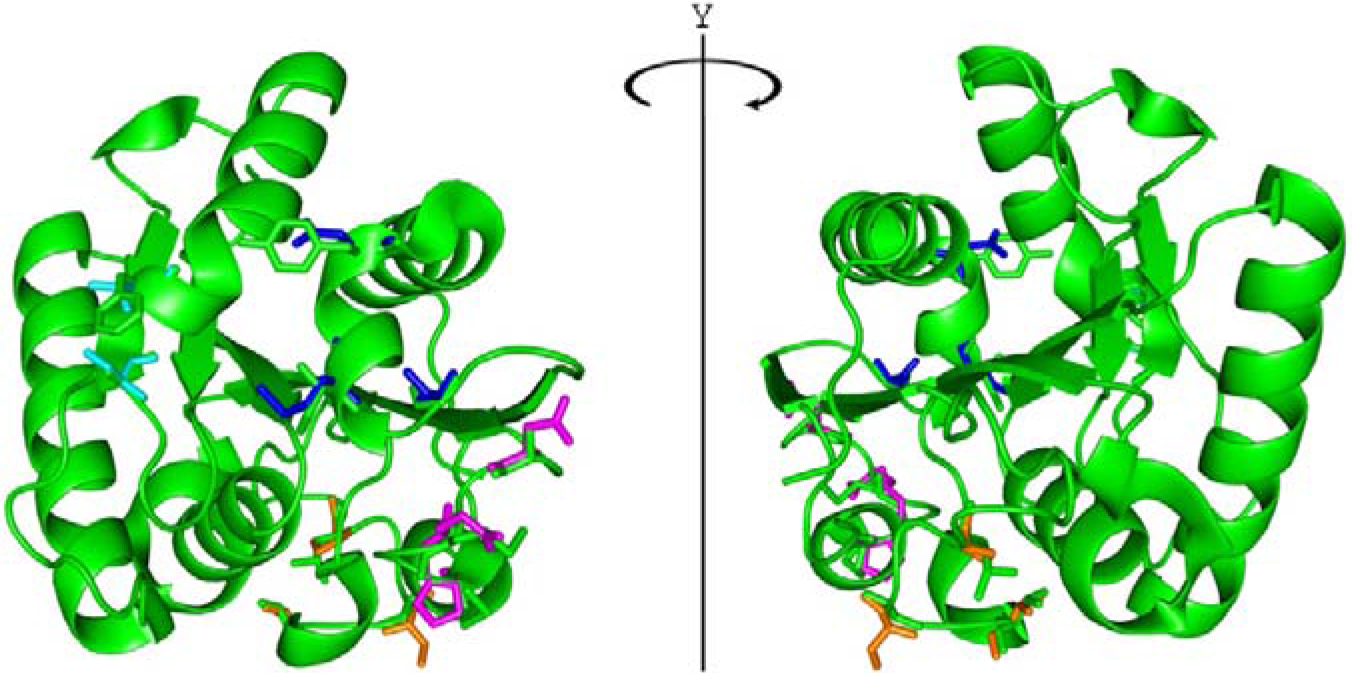
Positions of the mutations used in our study. Only the amino acids involved in mutation are shown as sticks. BSL(green), CM1(cyan), CM2(orange), CM3(magenta) and CM4(blue).

### Thermal unfolding

50 mM sodium phosphate buffer at pH 7.2 (PB) containing 50 μg/ml of protein was taken in a 1 cm path-length cuvette and heated from 25°C to 95°C at a ramping rate of 1°C/min. Change in ellipticity at 222 nm was monitored with respect to temperature by a JASCO J-815 spectropolarimeter. The temperature was controlled by Jasco peltier-type temperature controller (CDF-426s/15) during the experiment. The fraction of unfolded protein was calculated by using the following equation:

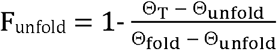

Where F_unfold_ is the fraction of protein which remains unfolded, Θ_T_ is the ellipticity of the protein solution at a given temperature, Θ_fold_ and Θ_unfold_ are the ellipticites of protein solution in folded and unfolded states respectively.

### Thermal aggregation

Aggregation kinetics, at a temperature, was evaluated by monitoring the scattering using Fluorolog 3-22 fluorimeter, fitted with peltier based cuvette holder for temperature control. Monochromators were set at 360 nm and slit width was set at 2 nm. Preheated (50°C) PB was taken in a fluorescence cuvette. Protein was added such that the final concentration becomes 50 μg/ml. Scattering was monitored for 10 min, immediately after addition of the protein.

### PNPB hydrolysis

10X substrate stock solution was prepared by micellizing 20 mM PNPB with 200 mM Triton X-100. The stock solution was diluted ten times by PB. 1 μg of lipase was added to 1 ml of substrate and the hydrolysis was followed by monitoring the absorbance at 405 nm at 25°C using a thermostated spectrophotometer for 2 min. Specific activity was calculated as the micromoles of PNPB hydrolyzed per minute per mg of lipase. The molar extinction coefficient of PNPB was taken as 1.711 mM^−1^.

### Susceptibility to proteolysis

For proteolysis assays, proteins (0.2 mg/ml in 20 mM Tris Cl pH 8.0 with 2 mM CaCl_2_) were incubated at 50°C with or without protease in a Bio-Rad thermal cycler. Proteolysis was checked against two non-specific proteases viz. Subtilisin A and thermolysin. Protease to lipase ratio of 1:50 (w/w) was used for proteolysis. Reactions were incubated for different time points (0, 20, 60, 120 min). Reactions were diluted 10 times with 50 mM sodium phosphate buffer (pH 7.2) and residual lipase activity was monitored by PNPB hydrolysis as mentioned above.

### Modelling protein structures

Protein structures were modelled by homology modelling using Modeller 9.14(24). At first, missing atoms and residues in wild type *Bacillus subtilis* lipase (BSL) structure (PDB ID: 1ISP) were modelled and the resulting model served as a template for modelling mutant structures. Sequences of different mutants were aligned with that of BSL by ClustalW using palm matrix. Hetero atoms and water molecules were ignored and energy minimization was avoided, as these reduce model quality. 10 models were created for each mutant and the model having the least DOPE score was selected. PyMol was used for visualization and PyMod 2.0 was used for running Modeller scripts(25, 26).

### Molecular dynamics simulation (MDS)

MDS was carried out using GROMACS-5.0.4 package(27) using AMBER03ws all-atom force-field (28) and TIP3P water model(29). Protonation states of amino acids were assigned as observed at pH 7. Protein was placed in the center of a rectangular box. The dimension of the box was such that there is at least 1 nm distance between any protein atom and the edge of the box. Box containing protein was solvated with pre-equilibrated water box and neutralized by replacing water molecule with the required number of Na^+^/Cl^−^ ions. 1000 steps of energy minimization were carried out using steepest descent algorithms to allow dispersion of solvent around the protein. During this step, position restraint (1000KJ/mol/nm^2^) was applied on heavy atoms of protein to avoid distortion. The system was equilibrated for 100 ps of NVT followed by 100 ps of NPT ensemble, during which temperature and pressure get equilibrated. The production run was carried out in NPT ensemble for 110 ns and analysis was performed on last 100 ns unless mentioned otherwise. Bond lengths were maintained by LINC algorithm(30), the temperature was maintained by V-rescale thermostat(31) and the pressure was maintained by Parrinello-Rahman barostat(32) throughout the MDS. Short range electrostatic and van der Waals interactions were truncated at 1.2 nm. Long-range electrostatic interactions were calculated by particle mess Edward method(33). 2 fs time step was used for calculation. Coordinates were saved at every 10 ps.

## Result

### Thermal melting of WT-BSL and mutants

Change in α-helix content of BSL and mutants (CM1 to 4), with respect to temperature, was monitor by CD at 222 nm (Fig.2). All the proteins undergo a two-step folding-unfolding transitions. Structural deformation of BSL started around 54°C and completed by 58°C with the midpoint of the transition (T_m_) around 56.3°C. All the mutants retained their structure at a higher temperature than the wild type. CM3 showed the highest T_m_ of 63.7°C. T_m_ of CM1, CM2 and CM4 were found to be 59.6, 60.1 and 62.4 °C respectively.

**Figure 2:**
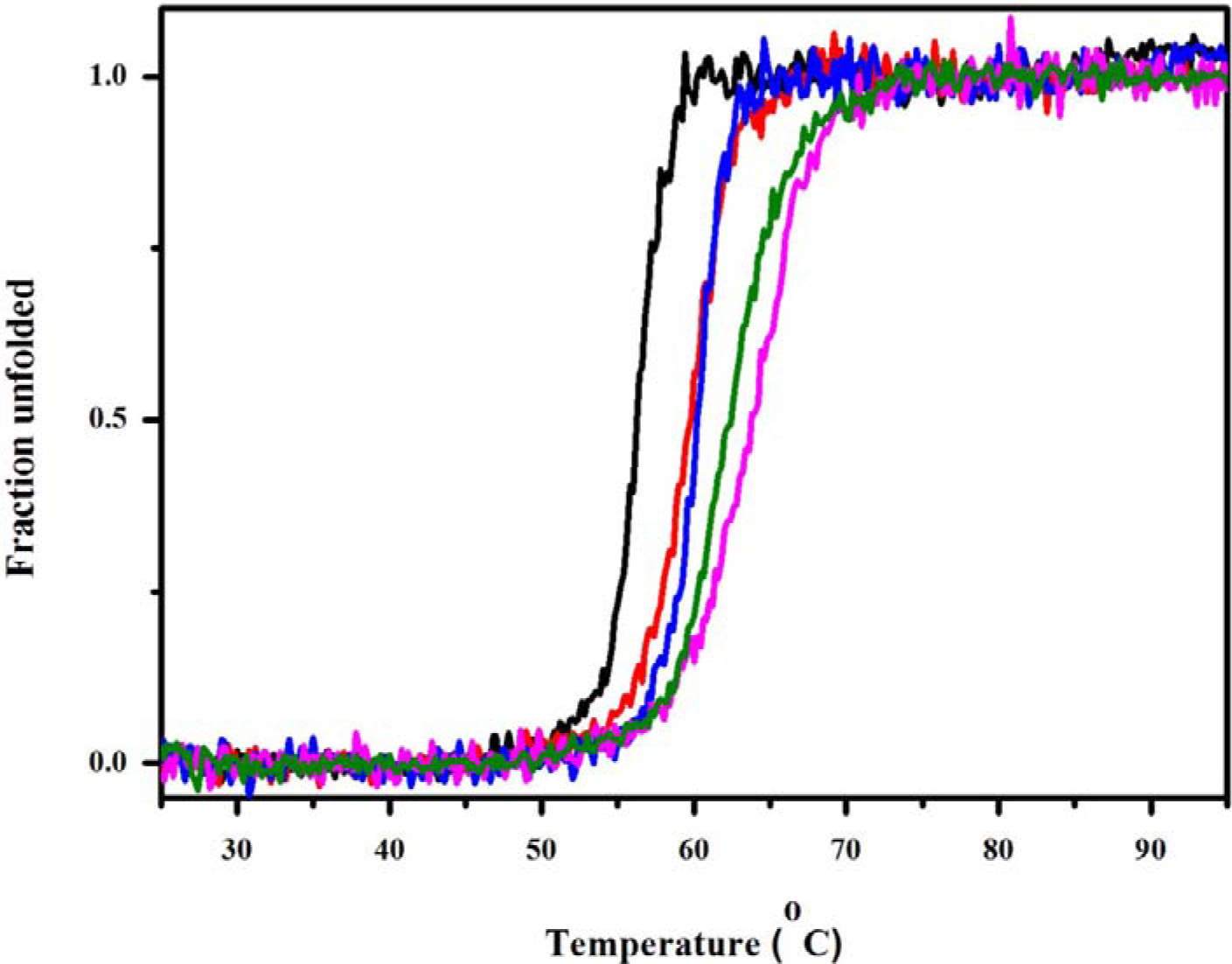
Thermal unfolding of BSL (black), CM1 (red), CM2 (blue), CM3 (magenta) and CM4 (green). The temperature was increased from 25°C to 95°C at a ramping rate of 1°C/min. The unfolding of protein was monitored as the change in ellipticity at 222 nm by CD.

### Aggregation kinetics

Aggregation kinetics of enzyme is crucial for its application in bioprocessing. Aggregation of BSL and mutants was measured by monitoring the scatter at 360 nm. As illustrated by Fig 3, the aggregation rate of all the mutants is far less than that of BSL with CM3 shows almost no sign of aggregation within the observation windows of 10 min.

**Figure 3:**
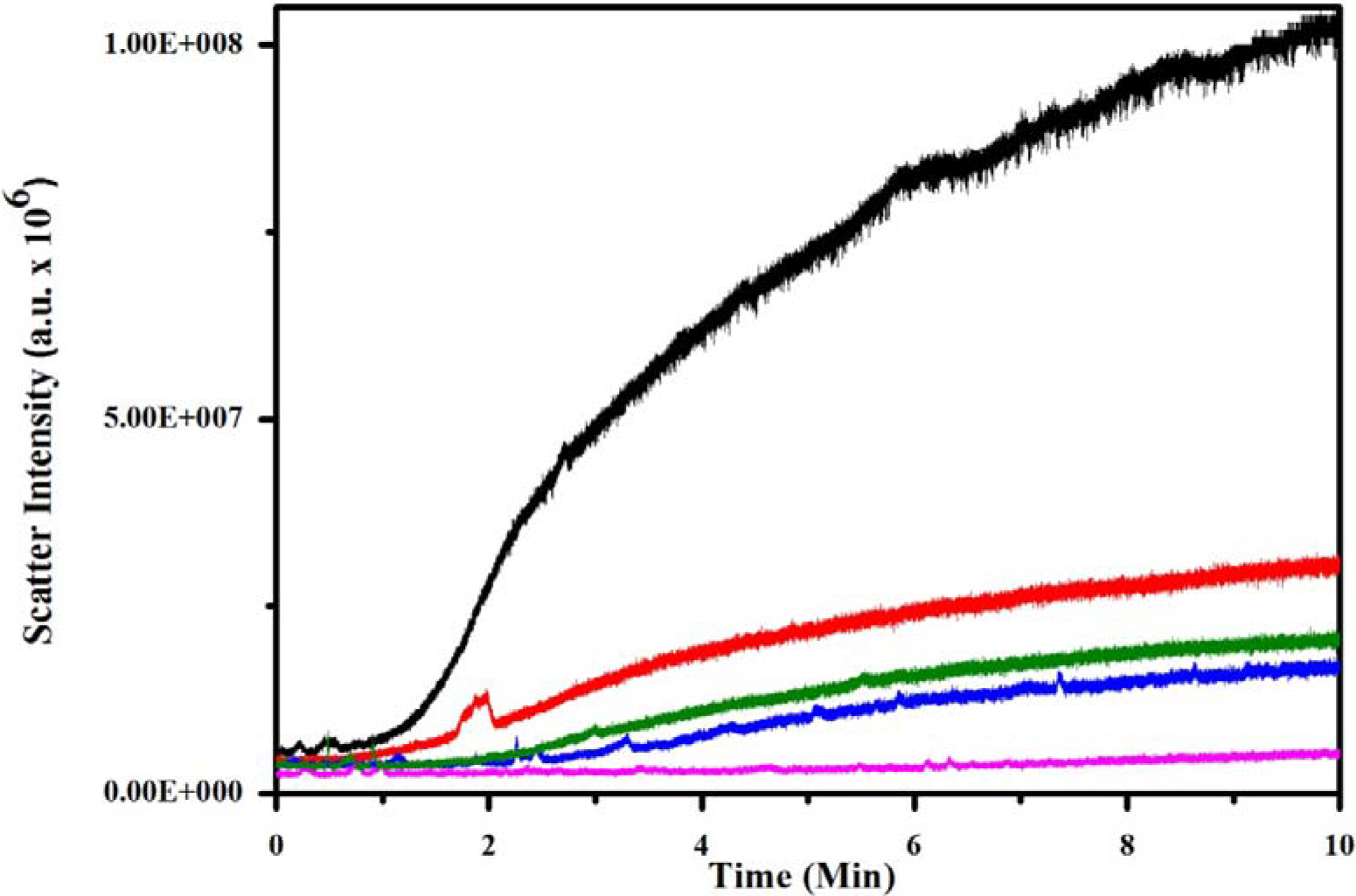
Aggregation kinetics of BSL (black), CM1 (red), CM2 (blue), CM3 (magenta) and CM4 (green). Concentrated protein solution was added to preheated buffer and scattering was measured at 360 nm for 10 min.

### Specific activity

Thermostable mutants usually show compromised activity. However, single mutations chosen for this study either had positive or neutral effect on the enzymatic activity. To study the effect of these mutations, upon combination, on the enzymatic activity, we monitored the hydrolysis of colorimetric substrate PNPB. PNPB and other *para*-nitrophenyl esters are frequently used as a substrate to quantify lipase activity due to their precise and colorimetric readout which can be monitored continuously. As illustrated in Fig 4, CM1 and CM4 didn’t show any change in activity compared to that of wild type BSL, whereas CM2 showed around 30 percent increase in activity while CM3 showed a remarkable 300 percent increase in specific activity with PNPB.

**Figure 4:**
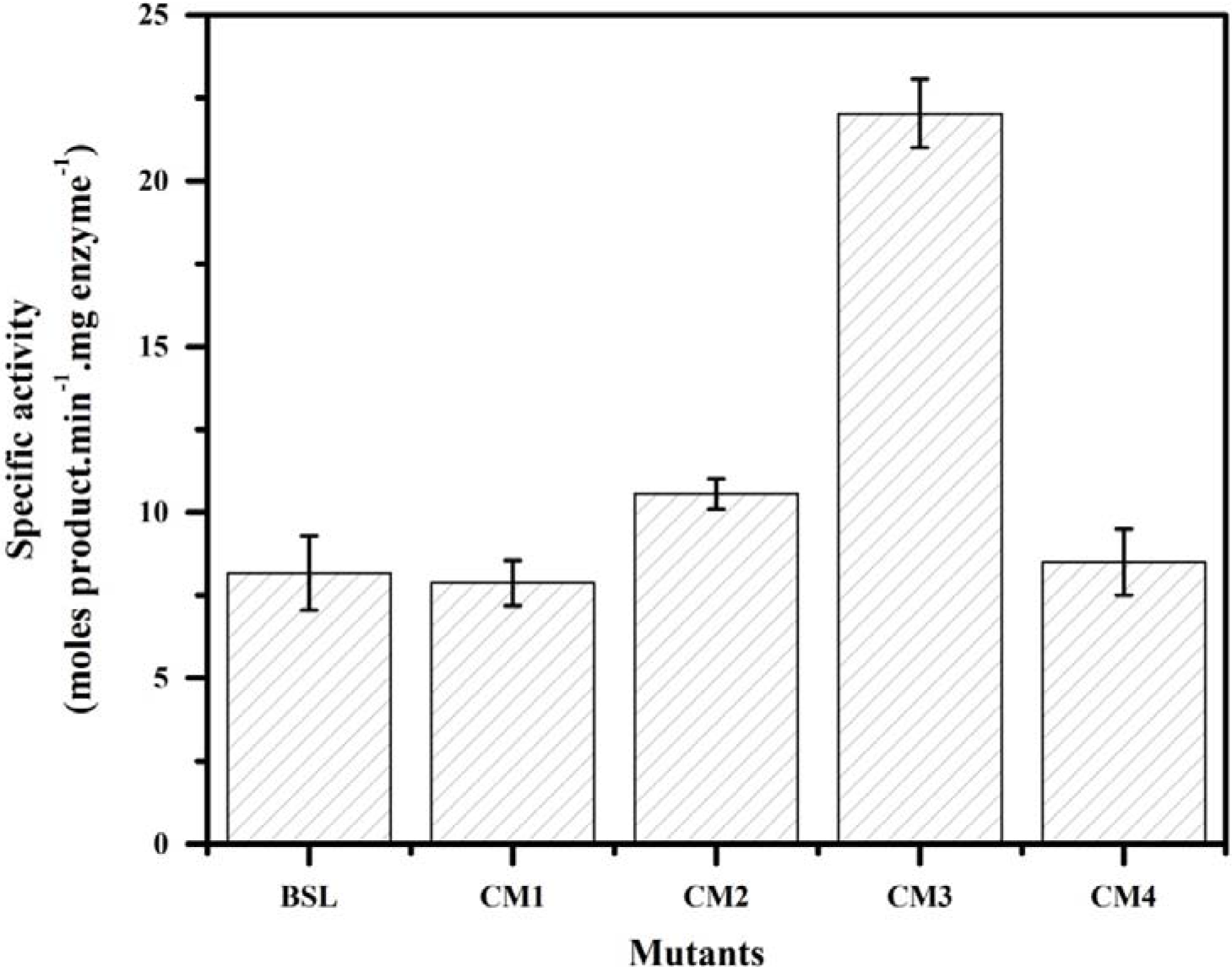
Specific activity of BSL and mutant lipases against PNPB. 1 μg of lipase was added to 1 ml of 2 mM PNPB and hydrolysis was monitored as the change in absorbance at 405 nm for 2 min.

**Figure 5:**
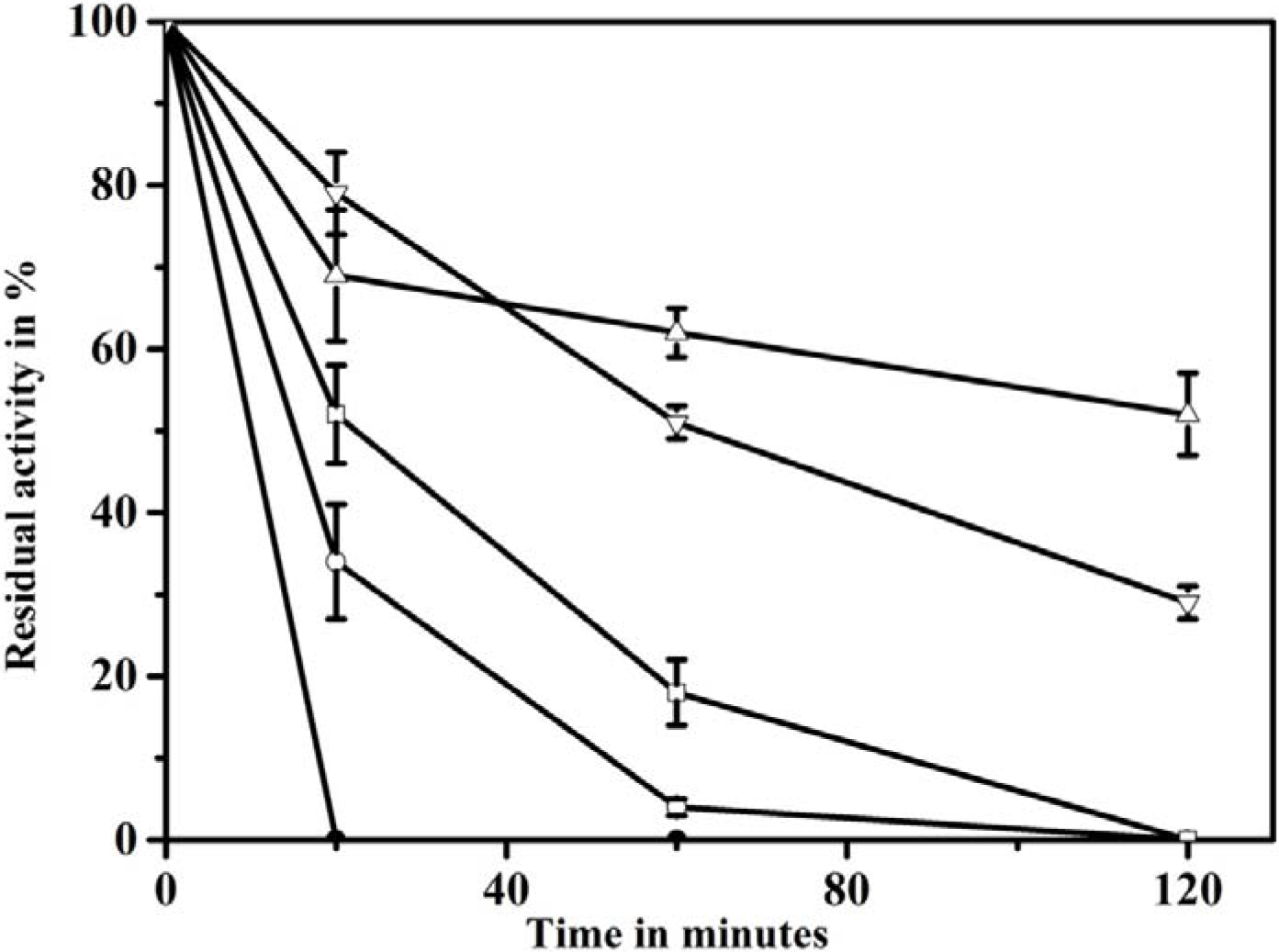
Proteolytic resistance of mutant lipases along with wild type against non-specific protease thermolysin. Wild type (Dark circle); CM1 (Open circle); CM2 (Open square); CM3 (Open triangle); CM4 (Inverse open triangle).

### Susceptibility to proteases

Susceptible to proteases is another indicator of stability of proteins. Proteolytic resistance of proteins was monitored by incubating proteins with non-specific proteases (thermolysin or subtilisin A) at 50°C. It was observed that wild type lipase had lost all its activity within 20 min while CM1 to CM2 showed different degrees of resistance to proteolysis against both proteases (Fig.5; see Supplementary Fig.2 for Subtilisin data). Wild type, CM1 and CM2 were completely inactivated by both the proteases within 120 min. CM4 retained nearly 20 and 30% activity after 120 min against subtilisin A and thermolysin respectively. CM3 retained more than 50% activity after 120 min against both the proteases. This is a remarkable improvement in proteolytic resistance considering the reaction temperature to be 50°C.

### Root mean square fluctuation (RMSF)

Mutations are known to alter protein stability and functionality by altering its dynamics(11, 12). RMSF calculated from MDS provides excellent measure of protein dynamics in the pico to nano second time scale. In order to understand the effect of mutations on protein backbone dynamics, we calculated the residue-wise RMSF of main chain heavy atoms (MC-RMSF) for BSL and its mutants (Fig 6). Excluding near terminal residues, maximum MC-RMSF was observed for residue near 119 in all the mutants. The magnitude of fluctuation in this region remains the same for all the mutants and BSL. Significant alterations in MC-RMSF values were observed to be concentrated into two regions, residue number 22 to 30 and 105 to 112 (will be referred to Region-I and Region-II respectively), irrespective of the positions of mutations. CM1, CM2 and CM3 mutant showed increased dynamics in the region-II, while CM4 showed increased dynamics in the Region-I compared to those of BSL. Amongst the mutants, CM3 showed the maximum increase in the dynamics which also correlate well with the observed enhancement in stability and activity.

**Figure 6:**
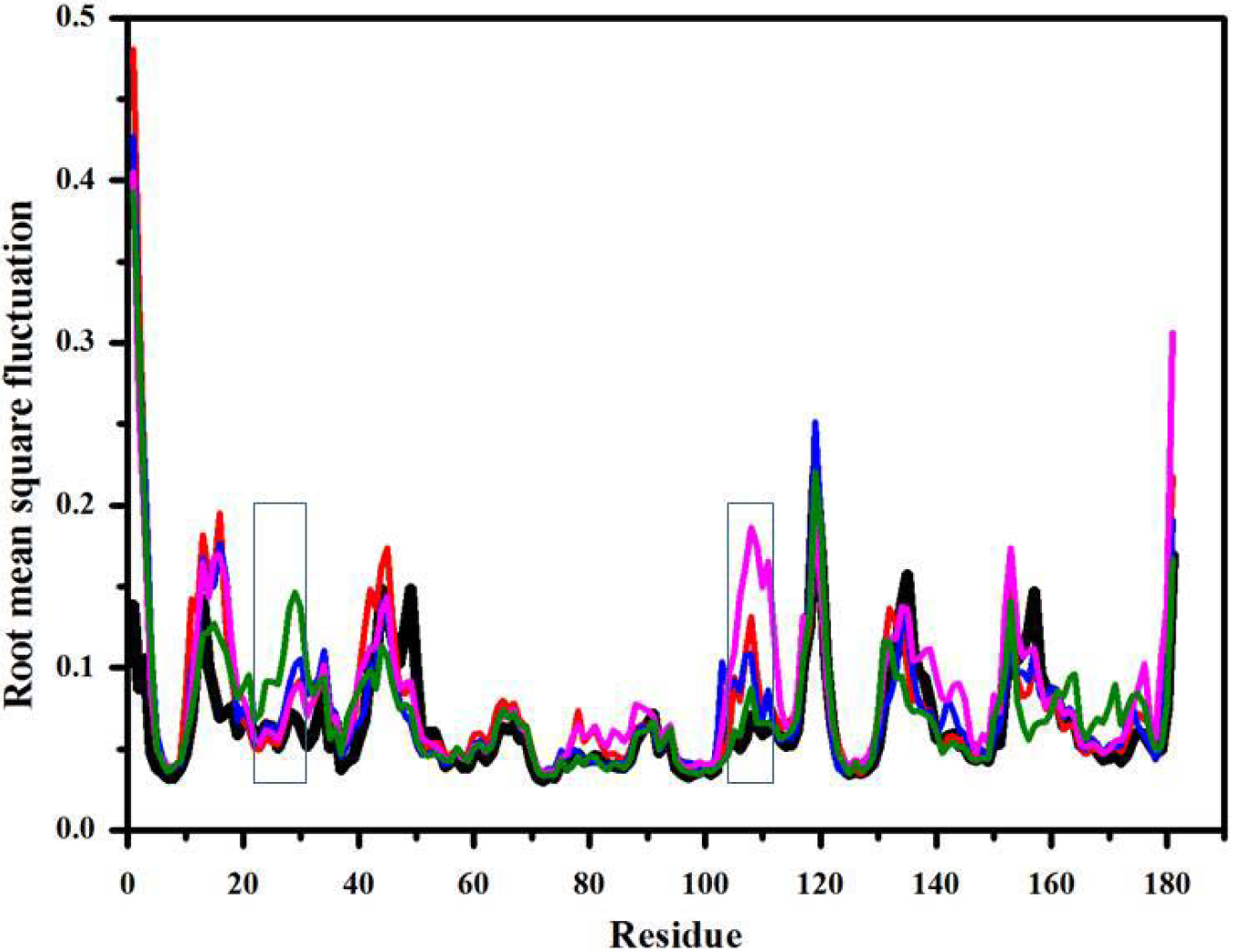
Root mean square fluctuation (RMSF) of atomic coordinates of main-chain heavy atoms of BSL (black), CM1 (red), CM2 (blue), CM3 (magenta) and CM4 (green). Average of root mean square deviation in atomic coordinates of non-hydrogen atoms were computed after performing least squares superposition of atomic coordinates.

### Squared order parameter (S^2^)

The squared order parameter (S^2^) obtained from NMR have been extensively used to study protein dynamics(34, 35). S^2^, calculated by using rotational correlation function from MDS, has been found to correlate well with that from NMR(36). Improvement of force-field along with affordable computation has made it a common practice to calculate S^2^ from MDS(37, 38). In the previous section, RMSF analysis of main chain atoms has shown that mutants have increased dynamics compared to BSL. In order to further investigate the effects of mutations, we calculated the S^2^ of amide N-H bond and Cα-Cβ bond (Fig 7). S^2^ of both N-H and Cα-Cβ provided an overall similar conclusion to each other and with that of RMSF analysis. All the mutants showed lower S^2^ (higher dynamics) compared to that of BSL with CM3 showing lowest S^2^ among all the mutants. Both the S^2^ showed an increase in dynamics near residue 150 which was insignificant in RMSF analysis. Due to this increase in dynamics, amino acids near 152 become the most dynamic region in mutants. Also, CM4 showed enhanced dynamics near residue 133.

**Figure 7:**
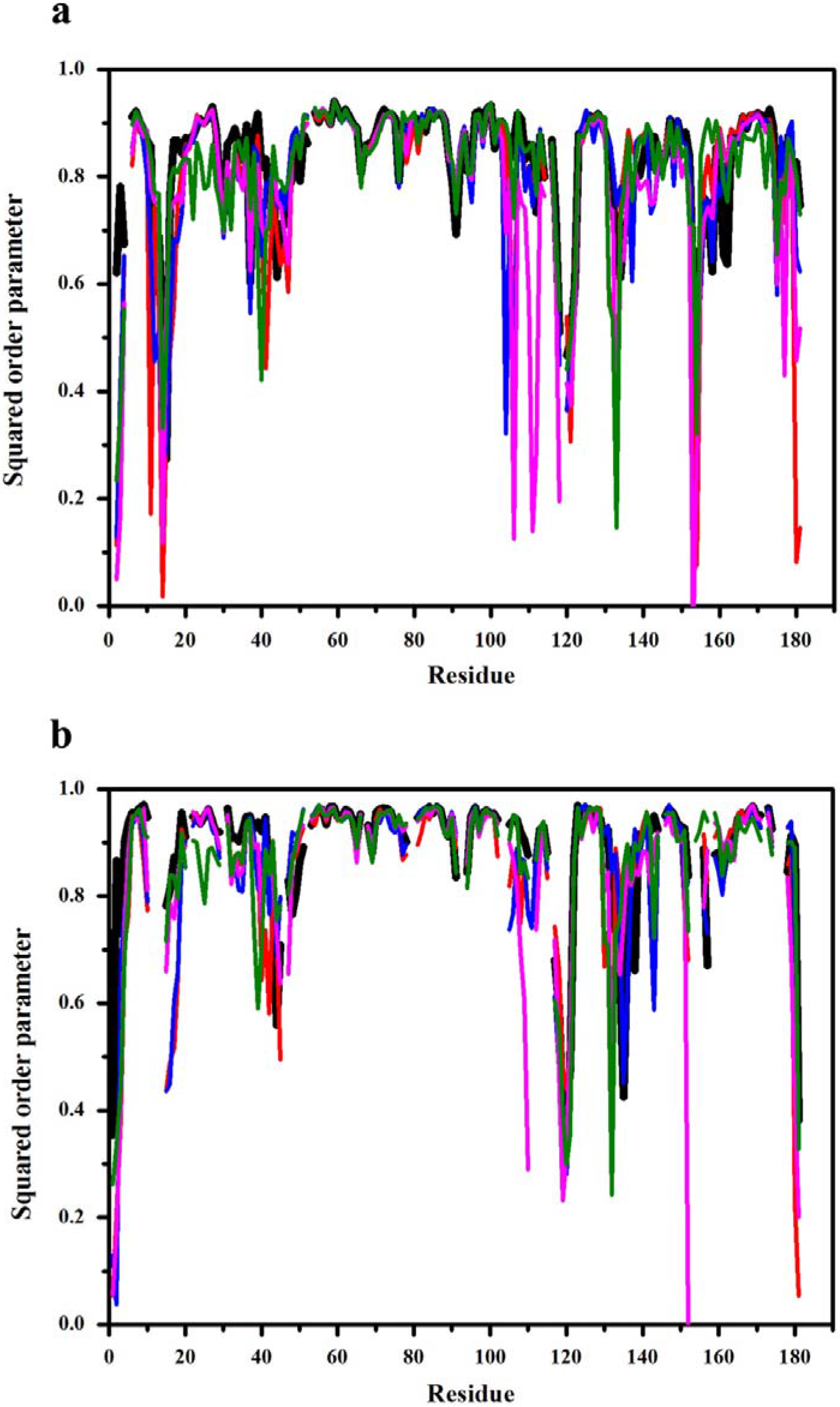
N-H (a) and Cα-Cβ (b) squared order parameter of BSL (black), CM1 (red), CM2 (blue), CM3 (magenta) and CM4 (green). Squared order parameter was calculated as the average rotational autocorrelation function of the respective bond vectors.

### Distance between mutation sites and site of increased dynamics

In order to investigate the effect of mutations on the local and global dynamics of the enzyme, we measured the average distance between the centre of mass of mutated amino acids and the centre of mass of the protein region which showed maximum alteration in dynamics. In case of CM1, CM2 and CM3 distances between the centre of mass of the region II and mutated amino acids were calculated whereas in case of CM4 distance between the center of mass of the region I and mutated amino acids was considered. Distances were calculated for each frame and averaged. In the case of CM1, CM3 and CM4 region showing altered dynamics were far from the mutations (Table 2).

**Table 2:**
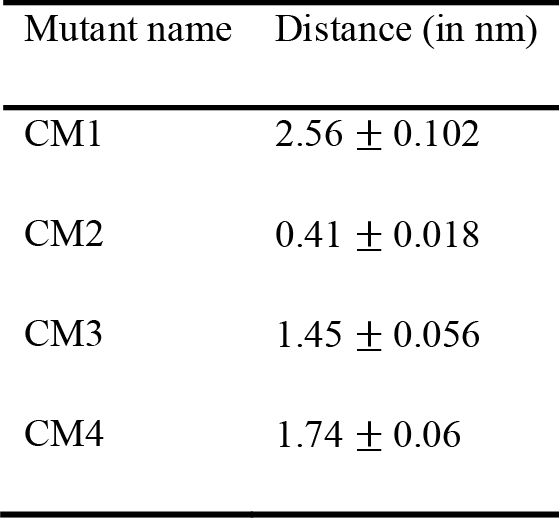
Distance between the mutation site and the site of altered dynamics. For CM1, CM2 and CM3 distance between the center of mass of region II and that of mutated amino acids were measured while for CM4 distance between the center of mass of region I and that of mutated amino acids was measured.

| Mutant name | Distance (in nm) |
| --- | --- |
| CM1 | $2.56 \pm 0.102$ |
| CM2 | $0.41 \pm 0.018$ |
| CM3 | $1.45 \pm 0.056$ |
| CM4 | $1.74 \pm 0.06$ |

### Distance between catalytic serine and region II

All the mutants investigated in this study showed increase in dynamics which is concentrated in to two regions. Out of these regions region II was closer to catalytic serine and showed increased dynamics for three out of four mutants. So we calculated the distance between center of mass of catalytic serine and that of region II in order to uncover any plausible correlation of this distance with that of enzymatic activity. Average distances and its standard deviations (SD) between catalytic serine and region II are listed in Table 3.

**Table 3:** Distance between the center of mass of the catalytic serine and that of the region II.

| Mutant name | Distance (in nm) |
| --- | --- |
| BSL | $1.26 \pm 0.044$ |
| CM1 | $1.27 \pm 0.063$ |
| CM2 | $1.32 \pm 0.044$ |
| CM3 | $1.31 \pm 0.080$ |
| CM4 | $1.16 \pm 0.044$ |

## Discussion

Effect of mutations on enzyme stability and activity is well established(39). Such alterations are mediated by structural change or change in dynamics. Several earlier studies have been conducted to explore the relationship between overall enzyme dynamics with its stability and activity(12, 40). Our study focuses the correlation between activity and stability with that of dynamics of different regions of enzyme. In this study we have created four variants of BSL (CM1 to 4) each of which contains two or three mutations. Each of these mutants contains mutations which are focused in a unique narrow region of BSL, mostly around a loop. These mutations along with their neighbours, taking together covers majority of BSL structure. We then evaluated the stability, activity and dynamics of each of these mutants and try to establish a correlation between them.

### Choice of mutation combination

Single mutants, obtained from the screening, show only marginal improvement in stability and activity. It is a common practice to combine mutations obtained from *in vitro* evolution to magnify their effect. Effect of such combination, particularly when they are nearby, may vary from synergistic to antagonistic and is difficult to predict. Very often such combination leads to deterioration of enzyme function rather than an enhancement. However, each of the combined mutant created for this study showed improved stability, though the improvement is not strictly synergistic. Use of different combinations might have resulted in better lipases. However, the aim of the study, which is to investigate the role of stabilizing mutation on local and non-local dynamics of enzyme, requires that the neighboring mutations should be combined. In this regard, one may argue that the neighborhood should be considered based on the vicinity in the structure rather than the sequence. Although justified, such an approach is difficult to execute as the structure of mutant may vary from that of wild type BSL, at least near the mutations, which would have led to a contradiction in neighborhood definition. Moreover, amino acids which occur nearby in the sequence are bound to be neighbours in the three-dimensional structure.

### Side chain RMSF doesn’t reflect local dynamics

While evaluating dynamics, we haven’t considered the RMSF of amino acids side chain. This is due to the fact that RMSF of amino acid side chain depends heavily on its type. For example, side chain of phenylalanine will always show more RMSF compared to alanine even though both of them experience similar dynamics (similar Cα-Cβ order parameter etc.) (42). Although mass weighted RMSF compensate for the above phenomenon to some extent, but the bias is still prevalent. This could give a false impression that the effect of mutations is restricted to its locality. To avoid such biasness, which would hinder the conclusion of this study, we excluded the RMSF of side chain from our analysis.

### Comparison of RMSF and S^2^

Protein dynamics observed by analyzing S^2^ gave a similar overall picture as that of RMSF analysis. The major difference between these two parameters was observed near residue 133 and 152 where CM3 and CM4 showed increased dynamics. Such minor variations are not surprising as the above two measures different modes of motion.

### Stability-dynamics correlation

Among all the mutants CM3 showed maximum improvement in stability. The same mutant also showed maximum enhancement in dynamics as illustrated by the increase in CM-RMSF and decrease in S^2^. Few similar observations have been reported earlier. Seewald *et al.* have shown that the stability of Streptococcal protein G is due to increased backbone flexibility(6). Talakad *et al*. created stable mutants of cytochrome p450, namely, 2B6 and 2B11 with increased dynamics (7). Dagan et al. stabilized a protein by increasing the dynamics of one of its loop (8). Similar relationship between activity and dynamics of enzymes have also been reported (5, 43–47). In our study CM1, CM2 and CM3 showed increase in dynamics in region II but not in region I while CM4 showed increased dynamics in region 1. This indicates that effect of stabilizing mutations are mostly confined to specific region of protein.

### Optimal dynamics varies between regions

All the four mutants studied here have shown increased MC-RMSF compared to that of BSL, which is concentrated primarily into two regions, excluding both the terminals. In BSL, both of these regions showed MC-RMSF value close to 0.05 nm which is very low compared to other part of the protein. However, the MC-RMSF of residue 118, 119 and 120, which showed the maximum MC-RMSF, is the same in all the mutants. In fact, no stabilizing mutation was observed in the vicinity of this region even though these positions were considered for mutagenesis in our previous work (23). This indicates that there could be an optimal value of dynamics. Exceeding this value may not further stabilize the protein or may even lead to destabilization by hindering with the necessary stabilizing interactions. Then the obvious question arises is “how the dynamics of other regions, whose MC-RMSF in BSL is of the order of 0.05 nm or less, affects the stability?” A long and continuous region is between 50 to 100 amino acid positions. The crystal structure of BSL indicates that most of the region, with little and unaltered dynamics (amino acid number 70 to 100), is buried in the protein interior while rest (amino acid number 50 to 67) of this region is part of the largest α-helix. Regions whose dynamics has increased significantly are completely surface exposed. This indicates that the value of optimum dynamics could be higher for surface exposed regions compared to that of buried region. This observation bridges two of the prior contrasting yet consistence observation which are 1) better core packing enhances protein stability(48, 49) and 2) increasing dynamics enhances protein stability(6).

### Protease susceptibility and dynamics

Like other stability parameters, resistance to protease susceptibility also is an indicator of protein stability. There is a common notion that higher dynamics increases protease susceptibility(50). However, in our study, mutants having higher dynamics showed less protease susceptibility. This can be explained as since proteases act mostly on denatured proteins, thermodynamic stability of the mutant is being translated as resistance to protease susceptibility.

### Activity-dynamics correlation

Like stability activity also showed a positive correlation with dynamics. This correlation was not observed for the dynamics of region I but observed only for the dynamics of region II, which is closer to the active site and makes several contacts with active site residues. CM3 which showed maximum activity also showed maximum SD in distance between region II and catalytic serine. BSL and CM4 which showed similar activity also showed similar SD in distance from catalytic serine. This observation further validates the common notion that activity of an enzyme correlates positively with its active site dynamics. However, CM1 which showed more SD has similar activity and CM2 which showed similar SD has better activity compared to BSL. In our study the mutations chosen for combination were primarily based on their stabilizing effect and therefore two out of four of our mutants didn’t show increase in activity. Understanding the relationship between activity and local dynamics requires different sets of mutations each contributing towards activity.

### Distance between mutation site and site of altered dynamics

In all the four mutants, significant enhancement in dynamics happened in certain locations. In the case of CM2, the mutation site is near the site of altered dynamics whereas in case of CM1, CM3 and CM4 these are far away in space. Such an observation, though compelling, is not out of line with the literature(40, 51–53). Such long distance effect is speculated to be through coordinated motion in protein or through alteration of hydrogen bond networks. Detail mechanism of such long distance effect requires further investigation.

## Conclusion

Our study is focused on interrelation between activity and stability of enzyme with that of its local dynamics. Our data suggests that there is optimal dynamics for stability and activity of enzymes. The optimal dynamics is different for different locations of the enzyme. Dynamics near the active site play crucial role in its activity. For stability, optimal dynamics of core region, as expected, is far less than that of outer regions. Stabilizing mutations altered the dynamics of different location towards its optimal value. Such alterations may happen in the vicinity of the mutation or far away from it. Outcome of this study will be helpful in understanding effect of mutation on enzyme stability and activity and will be applicable in engineering enzymes for more activity and stability.

## Author contribution

TRM has performed all the computational work and VK has performed all the experimental work presented in this submission. NMR has designed the study and prepared the manuscript.

## Acknowledgments

TRM and VK acknowledge the research fellowship received from CSIR. Authors acknowledge the generous computational support extended by Bioinformatics Research and Application Facility (BRAF) at CDAC, India. All the authors declare no conflict of interest with the work presented in this submission.

