## Supplementary Material for "Stability and dynamics interrelations in a Lipase: Mutational and MD simulations based investigations"

**For**

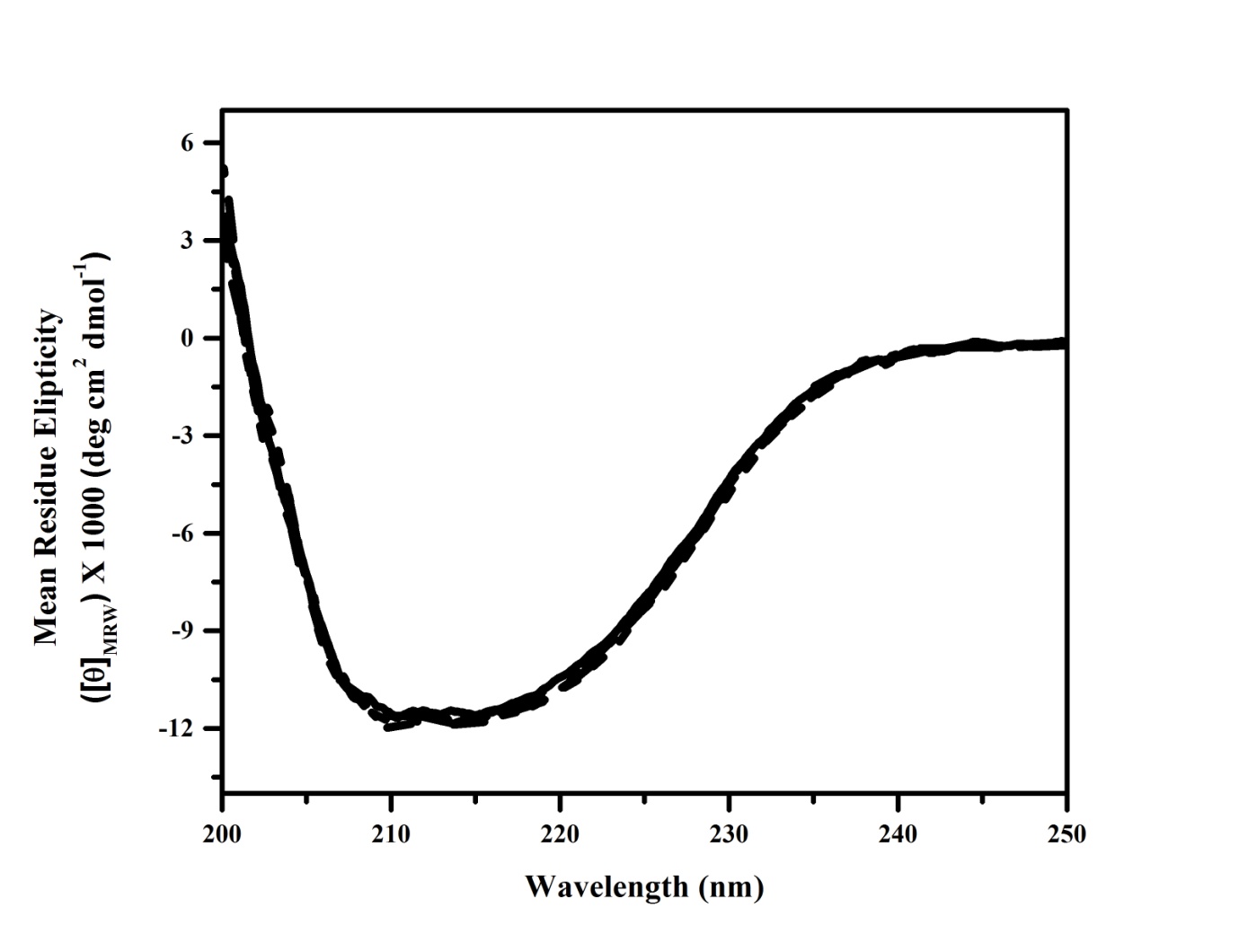

**Supplementary figure 1: Far UV CD spectra of BSL and mutants (CM1 to CM4).** Proteins were at 50 mM phosphate buffer and spectra were recorded at room temperature. All the mutants have the same secondary structure profile.

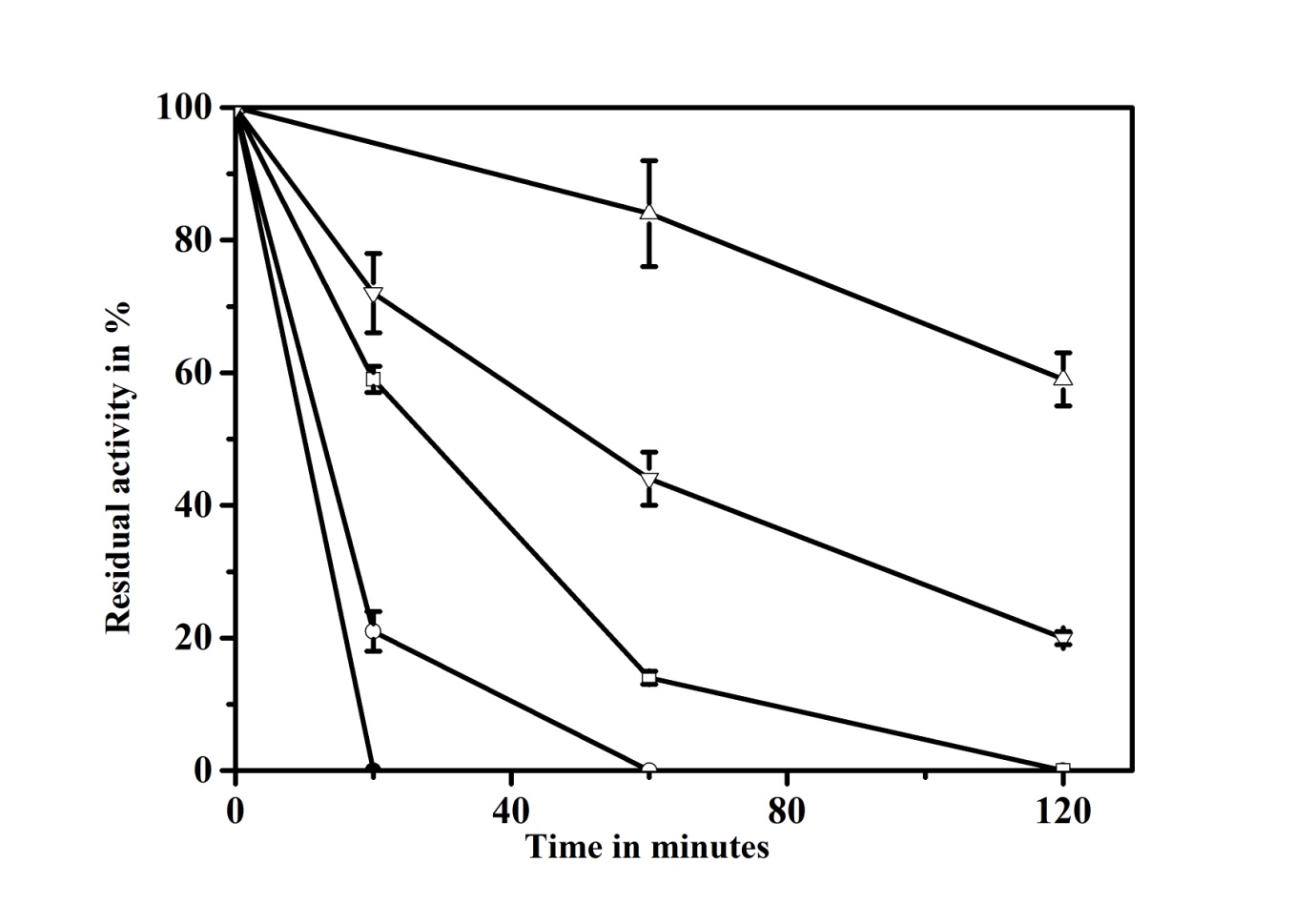

**Supplementary figure 2: Protease (subtilisin) susceptibility of BSL(closed circle), CM1(open circle), CM2(open square), CM3(open triangle) and CM4(inverted open triangle).** Proteins were incubated at 50⁰C with 50:1 (w/w) subtilisin and residual activity was monitored at 25⁰C. All the mutants showed less susceptibility to proteolysis.
